# Embryonic macrophages in brown adipose tissue are spatially associated with developing nerves

**DOI:** 10.1101/2025.01.16.632922

**Authors:** Inês Félix, Janni Ollikainen, Afra Haque, Joonas Karhula, Marko Salmi, Heli Jokela, Pia Rantakari

**Author notes:** Equal contribution (Inês Félix, Janni Ollikainen, Heli Jokela, Pia Rantakari).

## Abstract

Brown adipose tissue (BAT) is a specialized adipose tissue that produces heat through sympathetic nerve-dependent thermogenesis and influences glucose and triglyceride metabolism, making it a potential target for obesity-related diseases. In mice, interscapular BAT (iBAT) emerges on embryonic days (E)14.5-15.5, with extensive vessel and nerve networks still maturing postnatally. These developmental stages are critical for BAT, impacting long-term BAT function. In adult mice, macrophages regulate sympathetic innervation and thermogenesis. However, research on BAT macrophage characters, ontogeny, and functions in early life is lacking. Using single-cell genomics and proteomics, fate-mapping techniques, and 3D imaging, we analyzed iBAT macrophages during early development. We discovered the presence of fetal liver-derived macrophages as early as E15.5, which are later fully replaced by bone marrow-derived macrophages. Transcriptomics data highlight mononuclear phagocyte subpopulations, from which most express macrophage-associated genes (*Adgre1, Fcgr1, Csf1r*) and monocytic markers (*Ccr2*, *Ly6c2*) together. Embryonic macrophages interact closely with developing neural networks, while postnatal bone marrow-derived macrophages associate with blood vessels. Lack of fetal liver-derived macrophages disrupts nerve and vessel development and causes downregulation of genes related to neuronal diseases and protein digestion pathways. Our data suggest that fetal liver-derived macrophages, interacting with BAT’s developing sympathetic neural network, guide the development of tissue morphology. Understanding further the mechanism behind these events can unravel novel targets for BAT immunomodulation.

## Introduction

Brown adipose tissue (BAT) is a unique mammalian tissue that, by non-shivering thermogenesis, protects neonatal body temperature. Thermogenesis is stimulated mainly by the sympathetic nervous system (SNS) in response to cold exposure ^1^. The functional significance of BAT in newborn humans and small rodents has been widely recognized for some time. However, more recent research has unveiled that BAT also plays a significant role in the physiology of adult humans ^2–4^. The considerable capacity of BAT to dissipate energy and regulate triglyceride and glucose metabolism has stimulated interest in targeting this tissue therapeutically to fight obesity and improve glycemic control ^5–8^. In mice, interscapular BAT (iBAT) depots emerge at embryonic days (E)14.5-15.5, and by E17.5, brown adipocytes are already functionally differentiated ^9,10^. From birth to puberty, the extensive networks of vessels and nerves in iBAT are still maturing, while the tissue expands rapidly by primarily increasing brown adipocyte number ^11,12^. Brown adipocytes are fully developed in iBAT around 2 weeks after birth ^13,14^. BAT activity is barely identifiable in utero. However, iBAT activity rises right before birth, remaining robust until approximately two weeks of age in rodents ^14–16^.

Resident macrophages regulate healthy BAT’s thermogenic function in adult mice. Active adult BAT secretes CXCL14, a chemokine that promotes adaptive thermogenesis via macrophage recruitment ^17^. Furthermore, macrophages play a unique role in maintaining the sympathetic innervation of BAT, which is critical for energy expenditure homeostasis ^18^. Healthy adult BAT contains several distinct bone marrow-derived monocyte and macrophage subsets that intensely contribute to BAT macrophage maintenance and expansion in adult mice ^18,19^. However, it is well accepted that various tissue-resident macrophage populations are established before birth by embryonic precursors, either yolk sac macrophages or fetal liver monocytes, and are maintained in adult tissues independently of bone marrow hematopoiesis^20–24^. There is a tissue-specific combination of embryonic and adult macrophages in adult mice with distinct characteristics and functions. Nevertheless, the existence, origin, and functions of putative prenatal macrophages in BAT have not been studied.

iBAT is innervated by SNS, and the network of vessels is extensive, but very little is known about the factors affecting the development of the BAT microenvironment. In this study, we investigated the developmental dynamics of tissue-resident macrophages in prenatal and juvenile mouse BAT using high-dimensional single-cell transcriptomic and proteomic analyses combined with cell-fate mapping and macrophage-deficient mice. We revealed for the first time the existence of macrophage subpopulations already in embryonic iBAT under normal physiological conditions. Our data show that in the developing iBAT, the tissue-resident macrophages, with variable kinetics, are derived from fetal liver precursors. These embryonic macrophages are postnatally replaced by bone marrow monocyte-derived macrophages. Furthermore, we show that fetal liver-originated embryonic macrophages are nerve-associated during tissue development and that their deficiency associates with disrupted thermogenesis in postnatal mice.

## Results

### Macrophages show great complexity in developing BAT

To study and characterize the embryonic iBAT macrophages, we first performed flow cytometry analysis of pre (E15.5, E17.5) and postnatal day 0 (P0) murine iBAT. We pre-gated myeloid leukocytes (CD45^+^CD11b^+^) and analyzed the differential expression of the F4/80 cell membrane marker, which in previous studies has been shown to define F4/80^Int^ and F4/80^Hi^ macrophage populations originating from fetal liver and yolk sac, respectively ^25–32^. We showed the existence of macrophages in developing iBAT (Figure 1A; see gating strategy: Figure S1A). Interestingly, in pre and perinatal iBAT, the majority of iBAT macrophages are F4/80^Int^, and F4/80^Hi^ population is almost inexistent (Figure 1A), constituting less than 10 % of myeloid cells (Figure S1B). To understand the kinetics of macrophages during the development of the tissue after birth, we also collected iBAT from juveniles (P7 and P14) and adults (P56; 8-week-old). The F4/80^Hi^ macrophage population was not found in any of the time points studied (Figures 1A, 1B, and S1B). Interestingly, macrophage numbers were significantly increasing during development, alongside the growth of BAT, reaching the highest amount at P7 and then decreasing (Figure 1C).

**Figure 1.**
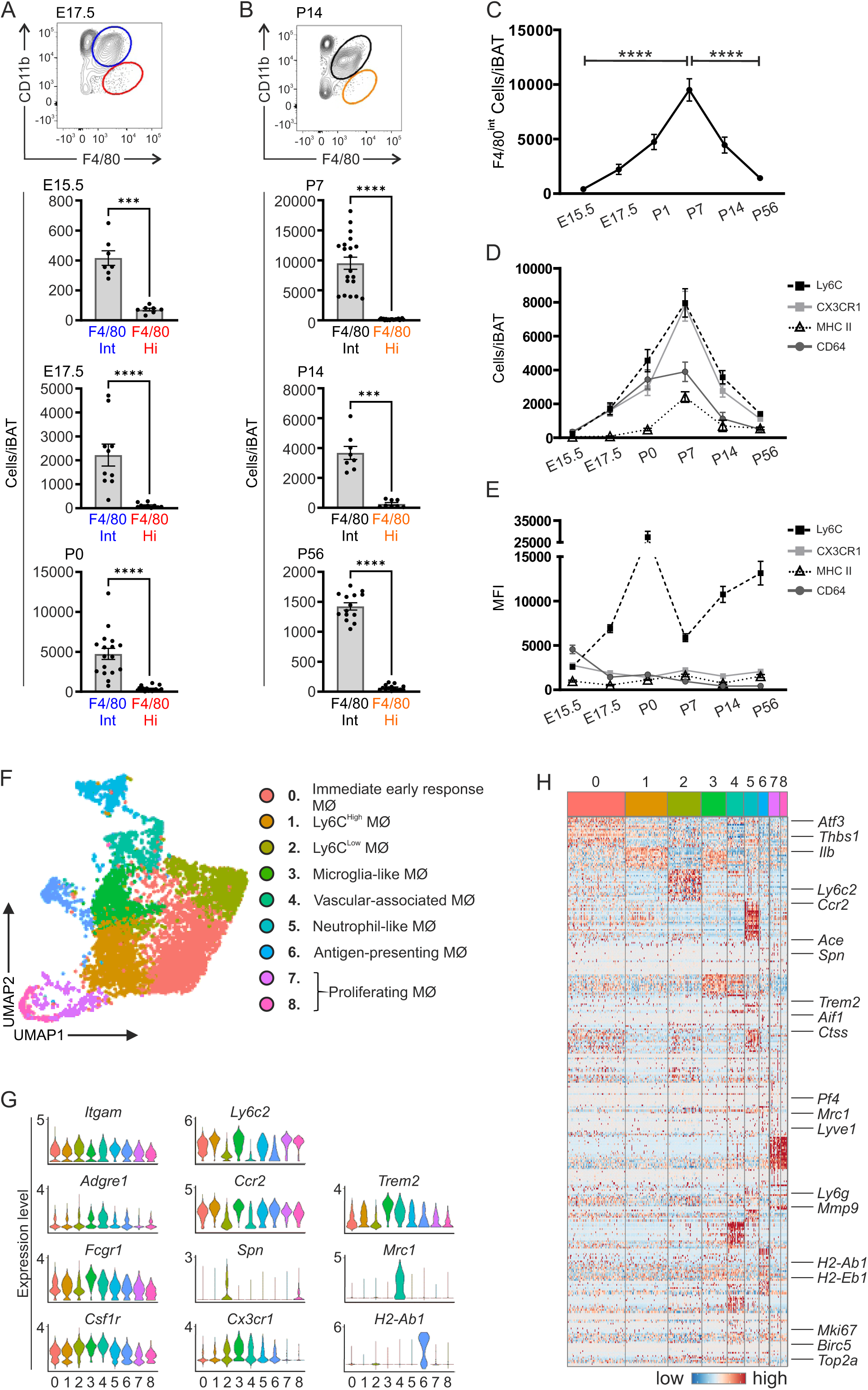
Single-cell analyses uncover heterogeneous signatures in brown adipose tissue macrophages during the development. (A and B) Representative flow cytometry plots and F4/80^Int^ and F4/80^Hi^ cell numbers per iBAT at indicated time points (n = 7-20/time point). (C) F4/80^Int^ cell numbers at indicated time points (n = 7-20/time point). (D) Cell amount of Ly6C, CX3CR1, MHC II, and CD64 positive cells at indicated time points. (E) Mean fluorescence intensity (MFI) of Ly6C, CX3CR1, MHC II, and CD64 from F4/80^Int^ cells at indicated time points. (F) UMAP visualization of iBAT macrophages expressing *Adgre1*, *Fcgr1*, and *Csf1r* (pooled data from E17.5, P0, and P7 mice). (G) Expression level as violin plots of markers identified in BAT macrophages. (H) Gene expression heatmap of markers identified in BAT macrophages.

When we further analyzed the F4/80^Int^ iBAT macrophage population, we found that there was a rise in the number of F4/80^Int^Ly6C^+^ cells that peaks at P7 of age, followed by a rapid decline towards adulthood (Figure 1D). However, the Ly6C expression in F4/80^Int^ macrophages was upregulated already earlier and was at its highest on P0 (mean fluorescent intensity, MFI; Figure 1E). Interestingly, the Ly6C expression level decreased from the P0 despite the expansion of cell numbers at the age of P7 (Figures 1D and 1E). Then again, Ly6C expression increased steadily from P14 onwards, although Ly6C^+^ cell numbers constantly decreased (Figures 1D and 1E). We then studied the expression of common macrophage markers CD64, CX3CR1, and MHC II (Figures 1D and S1C). Notable was that F4/80^Int^CX3CR1^+^ macrophage numbers followed the same kinetics as the F4/80 ^Int^Ly6C^+^ cells and increased until P7 and subsequently decreased through aging, even though the expression level stayed at a similar level throughout the development (Figure 1D and E). The expression level of MHC II and CD64 was identical throughout the development, although there was an increase in cell number at P7 (Figures 1D and 1E).

In adult mice, prolonged exposure to thermoneutral temperature (30 °C) leads to an increase in macrophage numbers ^33,34^. Therefore, we next investigated whether the early postnatal macrophages mimic the adult recruitment pattern. Pregnant female mice were acclimated to either thermoneutrality 30 °C or animal house standard room (semi-cold: 21 °C) temperature, and the delivered pups were kept at the corresponding temperatures during the weaning. Thermoneutrality seems to delay iBAT growth (Figure S1D). In juvenile iBAT, there was no alteration in the macrophage numbers associated with the different acclimatization temperature (Figure S1E), as opposed to what was previously described in adults ^33,34^. Interestingly, the elevated expression of CD11b observed under thermoneutral conditions coincides with the recruitment period of bone marrow-derived macrophages, suggesting that newly recruited iBAT macrophages at thermoneutrality exhibit more inflammatory profile ^20,24^ (Figure S1F). Together, these data prove the existence of macrophages already in prenatal iBAT. Most macrophages in developing iBAT have monocytic surface marker signatures, suggesting a monocytic progenitor origin.

### Most developing BAT macrophages express monocytic markers

To characterize the developing BAT leukocytes in detail, we performed single-cell RNA sequencing (scRNA-seq) of sorted live single CD45^+^ cells isolated from iBAT of E17.5 (n=8), P0 (n=6) and P7 (n=6) C57Bl/6N mice. Unsupervised clustering on integrated transcriptomic data showed various immune cell clusters within the iBAT leukocytes (Figure S1G). iBAT samples, regardless of time point, showed similar clustering, even though some clusters were mainly found from P0 and P7 samples (Figures S1H). To assess the identity of the cell type in each cluster, we examined the expression of canonical marker genes for different leukocyte cell types (Figure S1I). We confirmed the existence of lymphocytes (B and T cells), NK cells, mast cells, neutrophils, and basophils/eosinophils (Baso/eos). In addition, a clear cluster of macrophages (MØ) was one of the major cell types observed in early mouse iBAT (Figures S1G and S1I).

To further characterize the macrophages, we subclustered all *Adgre1*, *Fcgcr*, and *Csf1r* expressing cells, resulting in 9 different clusters in all time points (Figures 1F, S1J, and S1K). Most clusters had a monocytic signature with *Ly6c2* and *Ccr2* expression (Figure 1G and Table S1). Cluster 0 expressed typical macrophage markers (e.g., *Cd14*, *Il1b,* and *Dusp1*). This cluster also expressed *Thbs1*, *Atf3*, *Id3*, *and Nr4a1* involved in regulating cell death, including apoptotic processes (Figure 1H and Table 1). Cluster 1 showed a signature expression of the Ly6C^Hi^ population (e.g., *Ly6c2* and *Ccr2*), although having a clear macrophage core gene expression pattern (Figures 1G). Due to the low amount of differentially expressed genes when compared to the other clusters, the gene ontology (GO) enrichment analysis showed no specific function for this cluster. Cluster 2 was expressing genes typical for Ly6C^Low^ monocyte signature (e.g., *Plac8*, *Treml4*, *Ace*, *Cebpb*, and *Spn*; Figure 1G and Table S1) from which many are related to the regulation of stimulus (GO_0048583; Table 1). Interestingly, cluster 3 was enriched with genes related to activated microglia (e.g., *Trem2*, *Hexb*, *Aif1,* and *Ctss*; Figure 1G and Table S1), while cluster 4 had the expression profile of vascular-associated macrophages (e.g., *Pf4*, *Mrc1,* and *Lyve1*; Figure 1H). Both clusters shared functional terms general to macrophages, including immune system process and stress response (Table 1). Moreover, cluster 4 also expressed genes related to locomotion and endocytosis (Table 1 and Table S1). Clusters 7 and 8 presented a transcriptomic profile of the cell cycle and proliferating macrophages (e.g., *mKi67*, *Birc5,* and *Top2A*; Figure 1H). On the other hand, cluster 5 was identified as a neutrophil subset by the expression of *Ly6g*, *Retlng*, *Anax1*, and *Mmp9* (Figure 1H and Table S1), and cluster 6 as dendritic cells expressing genes such as *Cd209a*, *Ly6a*, *H2-Ab1*, *H2-Eb1,* and *Cd74*; Figure 1H and Table S1). Notable was that only cluster 6 expressed *H2-Ab1* and *H2-Eb1,* markers frequently detected in adult BAT macrophages and various other tissue macrophages ^18,19,35^ (Figure 1H and Table S1). Altogether, these results reveal the complexity of iBAT leukocyte content already during steady-state development and the diversity of macrophages. Furthermore, the monocytic signature of macrophages supports the idea of monocytic origin of the cells.

**Table 1.**
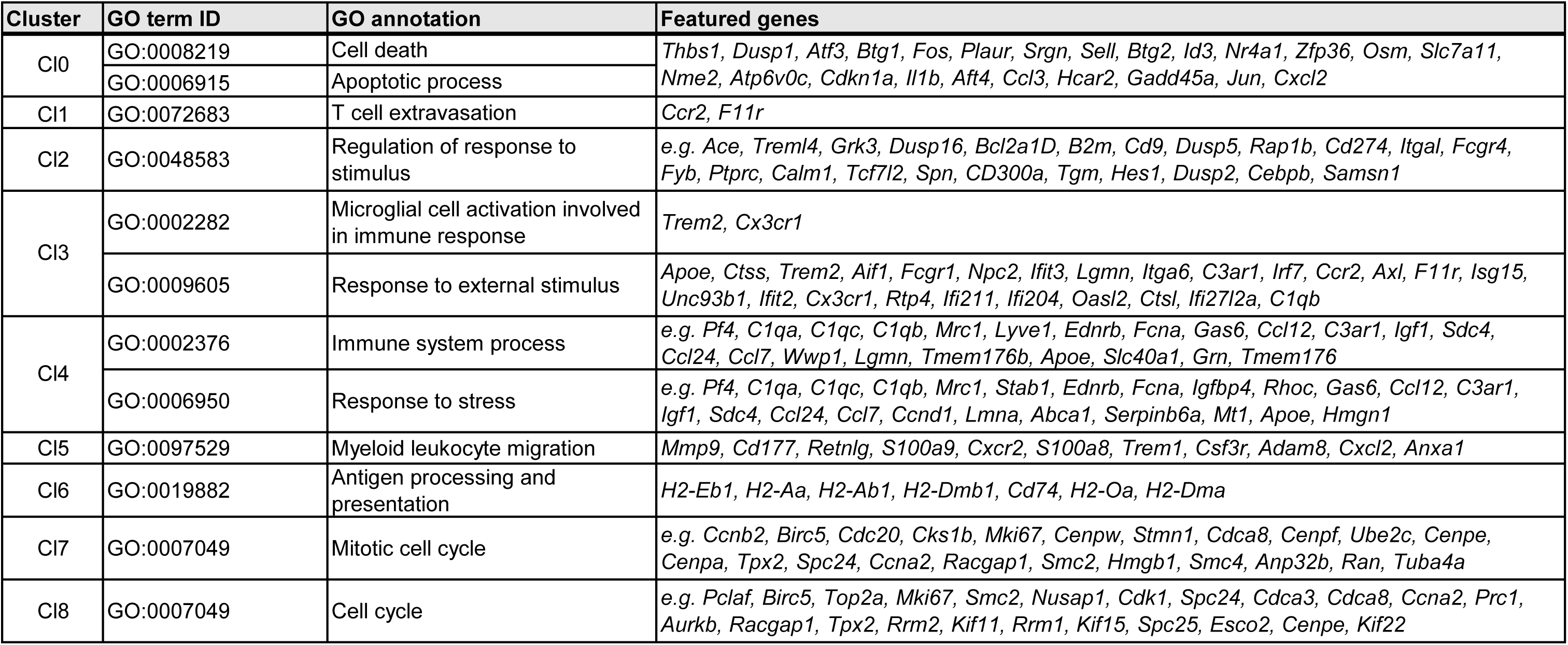
Gene ontology (GO) annotation of unsupervised clusters.

### Early BAT macrophages originating from the fetal liver progenitors are replaced with bone marrow-derived macrophages

In adult mice, the majority of BAT macrophages are derived from bone marrow ^19^. To study the unexamined early iBAT macrophage ontogeny, we used a combination of established macrophage reporter mice. Based on the observations in flow cytometry and scRNA-seq on the monocytic markers in iBAT macrophages, we first used *Ccr2^creERT^*^2^*; R26^TdTomato^* reporter mice, which allows the tracking of the Ly6C^Hi^ CCR2^+^ fetal liver-derived monocytes ^36–40^ (Figure 2A). At E17.5, iBAT macrophages were mostly labelled - TdTomato-positive (80 % of TdTomato^+^ cells were CD11b^+^F4/80^+^, Figures 2B, S2A, and S2B). However, already in iBAT of P14 *Ccr2^creERT^*^2^*; R26^TdTomato^* reporter mice, cell numbers and frequency of TdTomato^+^F4/80^+^ cells were diminished (Figures 2B and S2B). Notably, we no longer detected TdTomato-positive cells in iBAT of P35 (5-week-old) mice (data not shown), although the lung alveolar macrophages showed consistent TdTomato-positivity in all time points studied (Figure S2C).

**Figure 2.**
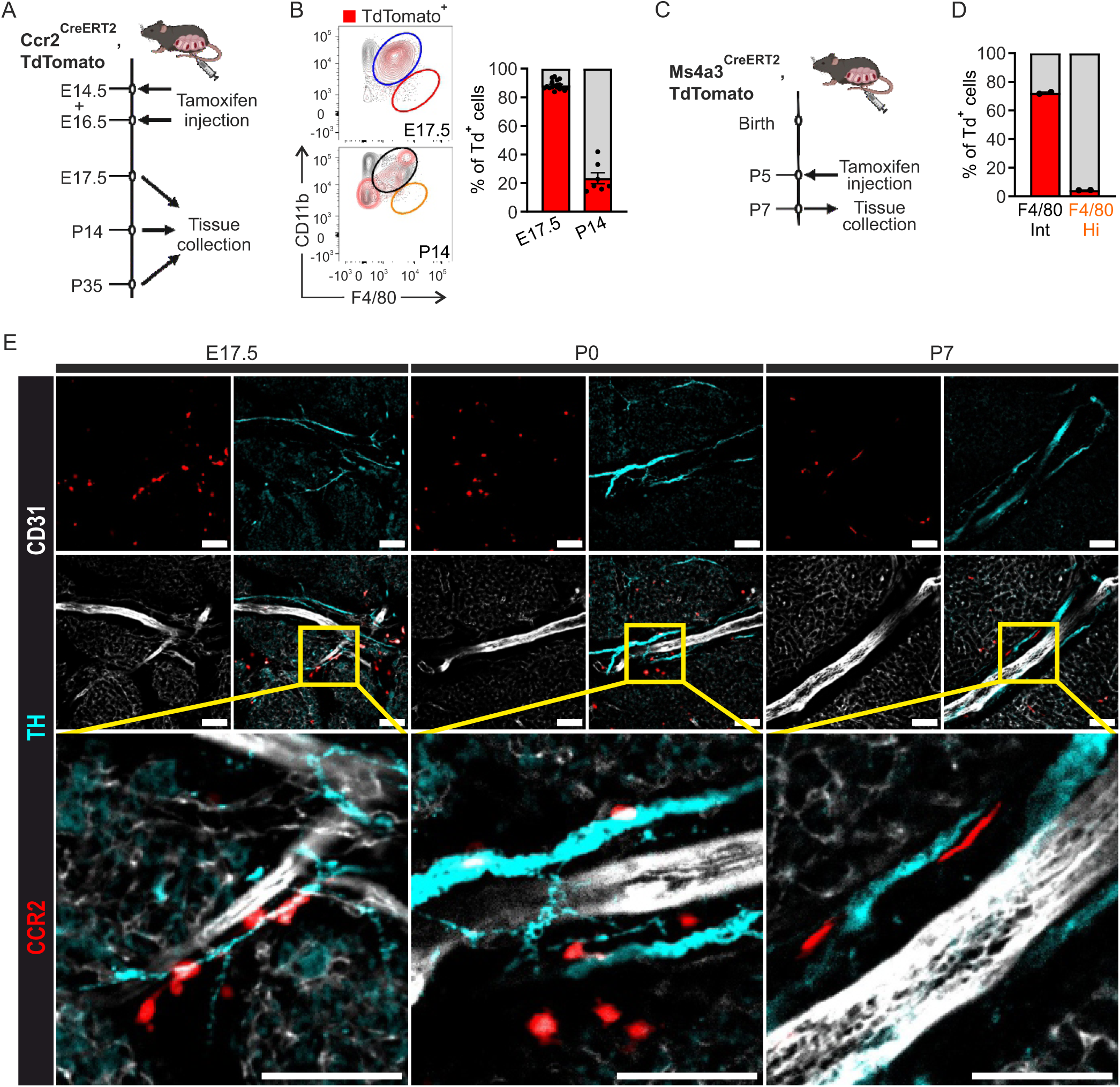
Fetal liver-derived macrophages dominate in early iBAT and interact closely with developing sympathetic neural networks. (A) Experimental setup of the induction scheme of *Ccr2^CreERT^*^2^*; TdTomato* mice. (B) Representative flow cytometry plots and TdTomato^+^ cell frequencies (positive cells backgated) of TdTomato^+^ and TdTomato^−^ in F4/80^Int^ cells (mean±SEM). (C) Experimental setup of the induction scheme of *Ms4a3^CreERT^*^2^*; TdTomato* mice. (D) TdTomato^+^ cell frequencies (positive cells backgated) of TdTomato^+^ and TdTomato^−^ in F4/80^Int^ and F4/80^Hi^ cells (mean±SEM). (E) Wholemount immunofluorescence staining of *Ccr2^CreERT^*^2^*; TdTomato* mice iBAT at E17.5, P0, and P7 with anti-tyrosine hydroxylase (TH; cyan), anti-CD31 (white), and anti-RFP (red) detecting TdTomato. Bar, representative of 50 µm.

Even though the yolk sac-derived F4/80^Hi^ population was almost entirely missing from iBAT (Figures 1A and 1B), we carried out a cell-fate mapping experiment to exclude the contribution of yolk sac-derived precursor to developing F4/80^Int^ iBAT macrophages. To that end, we crossed *Cx3cr1^CreER^*^2^ mice with *R26^EYFP^* mice and induced Cre with tamoxifen at E9.5 (Figure S2D). The labeling resulted only in a negligent amount of YFP^+^ cells in both F4/80^Int^ and F4/80^Hi^ iBAT macrophages of E17.5 embryos (Figure S2E). In contrast, brain microglia that originate solely from yolk sac-derived macrophages ^41^ showed around 50 % labeling (Figure S2F). These observations suggest that the yolk sac-origin macrophages are not required to fill or maintain the iBAT macrophage niche.

To understand how fast the bone marrow-derived macrophages infiltrate iBAT, we used the *Ms4a3^CreERT^*^2^*; R26^TdTomato^* model ^42^ with a single tamoxifen pulse five days after birth (Figure 2C). Two days after the tamoxifen injection (P7), blood monocytes were labeled at a substantial level (Figure S2G). Similarly, over 70 % of iBAT F4/80^Int^ macrophages were also TdTomato positive (Figures 2D and S2H), indicating that the influx of bone marrow monocytes in iBAT had already started. Altogether, the cell-fate mapping results show that early iBAT macrophages originate from the fetal liver precursors, while yolk sac-derived cells scarcely contribute to the tissue-resident iBAT macrophages. However, the iBAT macrophages of embryonic origin diminish already at two weeks of age and are completely replaced by bone marrow-derived monocytes at five weeks of age.

### Fetal liver-derived macrophages interact spacially with the developing nerve network in iBAT

To reveal the spatial localization of fetal liver-derived macrophages in developing iBAT, we performed immunofluorescence staining of optically cleared wholemount iBAT from E17.5 *Ccr2^creERT^*^2^*; R26^TdTomato^* mice. iBAT is highly vascularized, and macrophages are frequently associated with blood vessels in many tissues ^26,43,44^. Thus, we first studied whether the macrophages are vessel-associated during development by staining iBAT endothelial cells with anti-mouse CD31 and CCR2-positive macrophages with an anti-red fluorescent protein (RFP), detecting TdTomato. Interestingly, no apparent contact with the vasculature and TdTomato^+^ macrophages was seen, but instead, iBAT macrophages were forming clusters distributed throughout the tissue stroma (Figure S2I). In addition to being extensively vascularized, BAT features a dense network of sympathetic nerves, which regulate thermogenesis ^45,46^. To investigate whether the macrophage clusters interact with this intricate nerve network, we stained E17.5 iBAT with anti-CD31, anti-RFP, and a sympathetic nerve cell marker, tyrosine hydroxylase (TH). We observed a clear interaction between embryonic iBAT macrophages and developing nerves at E17.5, P0 and P7 (Figure 2E). TdTomato-positive macrophages were in contact with the nerves and formed a pear-like continuing structure following the nerves (Figure 2E). Taken together, these results indicate a possible role of fetal liver-derived macrophages in the development of nerve networks in iBAT.

### Prenatal macrophages deficiency correlates with morphological alterations in iBAT development

ScRNA-seq and fate-mapping analyses provided valuable phenotypic information from the heterogeneity and ontogeny of the BAT resident macrophages in homeostasis. To further study the functions of prenatal macrophage in BAT tissue responses, we evaluated the iBAT of E17.5 and P7 *Plvap^−/−^* mice, which show a reduction of fetal liver-derived macrophages but normal yolk sac and bone marrow-derived macrophages in most tissues ^22^. Flow cytometric analysis of E17.5 iBAT confirmed a significant decrease in the cell numbers of the F4/80^Int^ population (fetal liver-derived) but not in the very small yolk sac-derived F4/80^Hi^ population (Figure 3A). In adults, macrophage populations exhibit distinct F4/80 expression profiles based on their origin: both yolk sac- and fetal liver-derived macrophages upregulate F4/80 to become F4/80^Hi^, whereas bone marrow-derived resident macrophages are F4/80^Int^ ^30,47^. At P7, the cell numbers of the F4/80^Int^ population in *Plvap^−/−^*mice are already comparable to WT, most likely because of normal infiltration by bone marrow-derived macrophages (Figure S3A). Furthermore, at P7, when the F4/80 expression level of embryonic cells is still in transition, there is a tendency of decrease in the small population of embryonic-derived F4/80^Hi^ macrophages in iBAT of *Plvap^−/−^* mice (Figure S3A).

**Figure 3.**
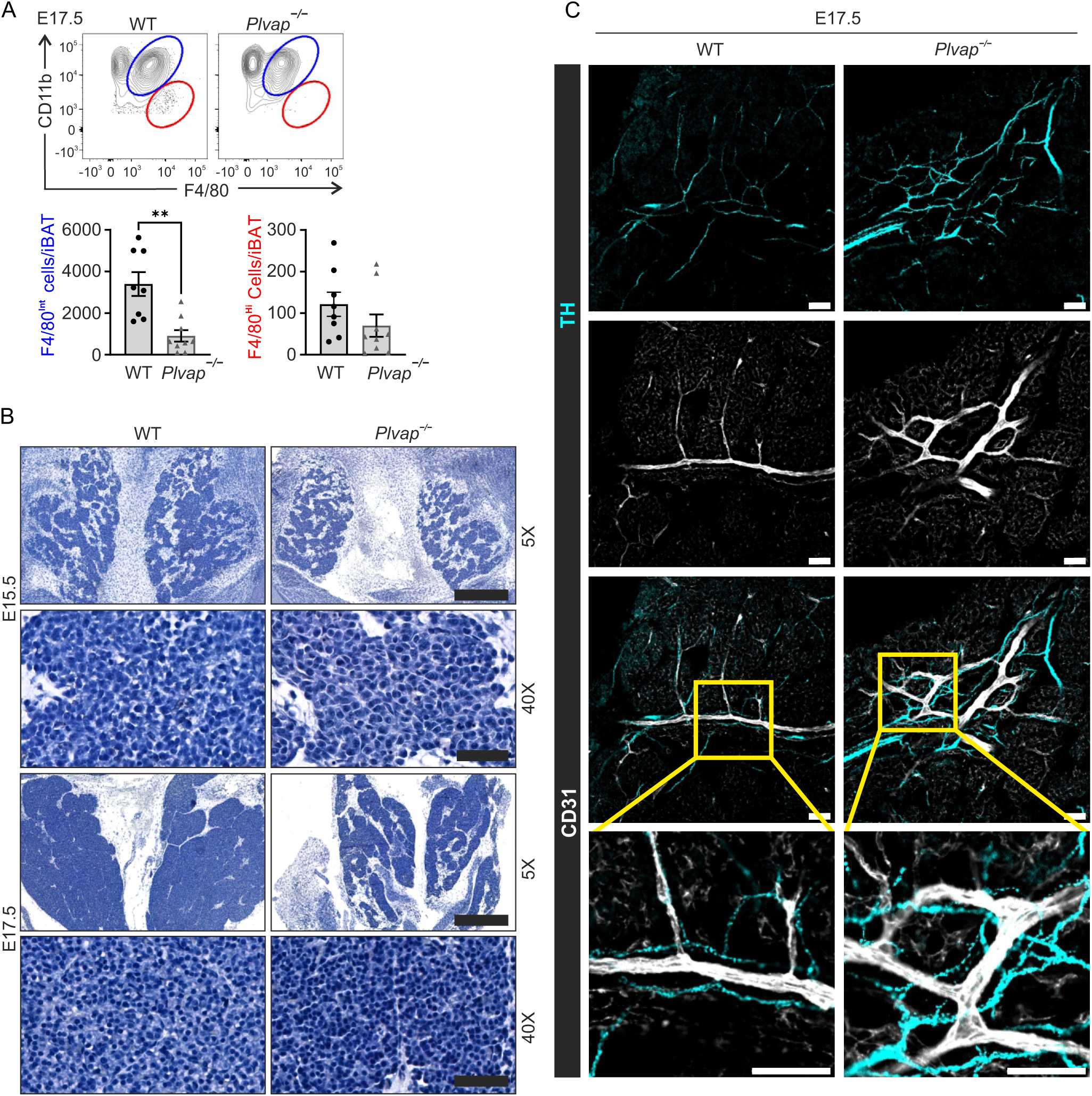
Defect in fetal liver-derived macrophages associates with a delay in embryonic iBAT development. (A) Representative flow cytometry plots and cell numbers of F4/80^Int^ and F4/80^Hi^ macrophages per iBAT of *Plvap^−/−^* and WT mice at E17.5 (n = 8-9). Mean±SEM. (B) PAS stained iBAT histology in *Plvap^−/−^* and WT mice at E15.5 and E17.5. Scale bars are 50 µm in overview (5X) and 500 µm in 40X images. (C) Wholemount immunofluorescence staining of *Plvap^−/−^* and WT mice iBAT at E17.5 with anti-TH (cyan) and anti-CD31 (white). Bar, representative of 50 µm.

Next, we studied whether the deficiency of fetal liver-derived macrophages affected iBAT morphology. Histologic analyses of prenatal *Plvap^−/−^*iBAT revealed delayed development, with islands of brown adipocytes failing to merge in *Plvap^−/−^* at E17.5, in contrast to the merged islands in WT littermates (Figure 3B). However, in postnatal mice, when bone marrow-derived macrophages are already present, relative iBAT weight and overall tissue architecture, with multilocular adipocytes and vasculature, was comparable with WT littermates (Figures S3B and S3C). Imaging of TH-positive nerves and CD31-positive blood vessels, however, showed that the lack of fetal liver-derived macrophages seemed to impair the sympathetic nervous network development and vasculature, as we detected both TH and CD31 signals in pre- and postnatal iBAT of *Plvap^−/−^*mice (Figures 3C and S3D).

### Macrophage deficiency in developing BAT associates with dysfunctional BAT in newborn and juvenile mice

We next assessed how the deficiency of fetal liver-derived macrophages during embryonic tissue developmental period affected BAT function at its most active time, P14 ^16^. We isolated total RNA from iBAT of P14 *Plvap^−/−^* and WT littermate mice to conduct bulk RNA sequencing. In the principal component analysis, the *Plvap^−/−^* samples were clearly distinguished from the WT group (Figure S4A). Furthermore, the effect of macrophage deficiency was evident, as 792 genes were uniquely expressed in *Plvap^−/−^* and 420 in WT iBAT (Figure 4A). Heatmap for the expression values of *Plvap^−/−^* and WT revealed accurate group assignment, indicating the presence of uniquely and significantly different expression patterns for the differentially expressed genes (DEGs; Figure 4B). The DEG analysis of *Plvap^−/−^* iBAT revealed significant downregulation of genes related to BAT specific functions, such as thermogenesis (e.g. *Cox6b1*, *Cox6b2*, *Uqcr11*, *Atp5g3*, and several *Nduf* genes), PPAR signaling (e.g. *Acaa1b*, *Pltp*, *Fads2*, *Hmgcs2*, and *Slc27a2*), and adipogenesis (e.g. *Col6a2*, *Hrasls*, *Gdf6*, and *Nxn*). Furthermore, genes related to neuronal activity were significantly downregulated, e.g. *Mbp*, *Matn2*, and *Prx* (Figure 4C). On the other hand, genes related to immune functions were found to be upregulated in *Plvap^−/−^*iBAT (e.g., *Il6* and *Ly6i*). Interestingly, IL6, in addition to macrophages, can also be produced by brown adipocytes upon β-adrenergic activation and act as a thermogenic cytokine ^48^. To understand the biological function of the differentially expressed genes, we performed pathway enrichment analysis (Figures 4D and S4B). As expected, various upregulated genes were associated with pathways and GO annotations related to immune functions, such as Il-17 signaling and antigen processing and presentation (Figures S4B and S4C). Notably, the functional enrichment analyses of the top 196 downregulated genes showed that many DEGs were involved in thermogenesis and the PPAR signaling pathway (Figure 4D). In addition, both KEGG and GO analyses revealed an enrichment of downregulated genes in several extracellular matrix (ECM) interactions. Recently, ECM interactions have been shown to influence brown adipocyte formation^49^. In addition, KEGG analysis revealed significant downregulation of pathways related to neuronal diseases (Parkinson and Alzheimer) and protein digestion and absorption. All in all, these results indicate that the deficiency of fetal liver-derived macrophages during iBAT development may impact early BAT function.

**Figure 4.**
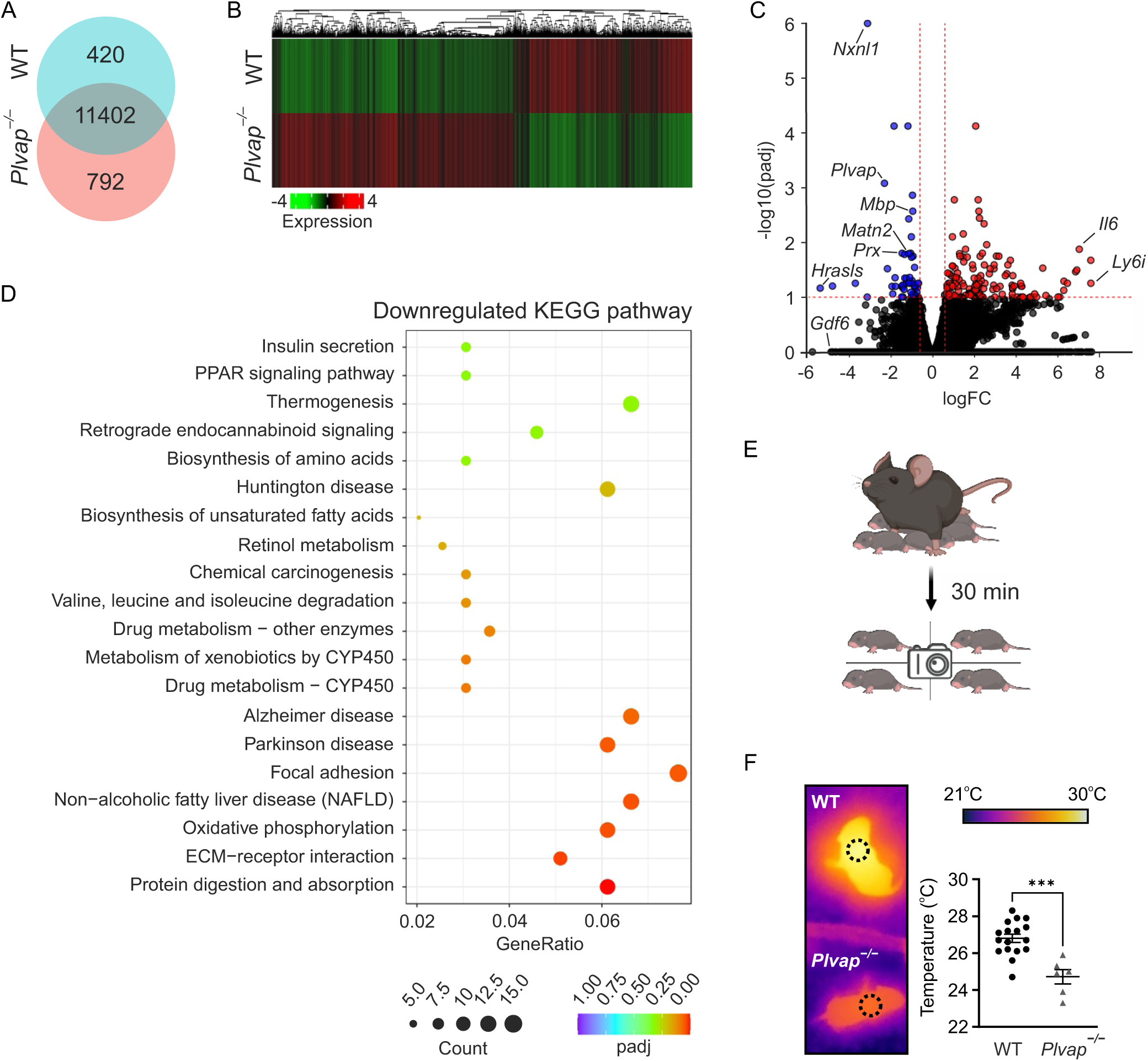
Macrophage deficiency during development correlates to dysfunction of early iBAT. (A) VENN diagram of shared and unique transcripts in P14 iBAT of *Plvap^−/−^* and WT mice determined by Bulk RNA sequencing (n = 4). (B) Heatmap of the differentially expressed genes in *Plvap^−/−^* and WT iBAT bulk RNA sequencing data. (C) Volcano plot of differentially expressed genes in iBAT of *Plvap^−/−^*. (D) KEGG enrichment analysis of the top 20 downregulated pathways in iBAT of *Plvap^−/−^*. (E) Experimental setup for infrared thermography analysis from P5 *Plvap^−/−^* pups. (F) Representative thermographs and interscapular iBAT temperature of *Plvap^−/−^* and WT littermate P5 pups (n = 6-16).

Considering the vital role of thermogenesis for newborn mice survival, we next wanted to test if the deficiency of fetal liver-derived macrophages affects the postnatal thermogenesis of *Plvap^−/−^* on the first days after birth. We used an experimental setup where we separated the pups from their mother for 30 min at room temperature and employed infrared thermography to quantify iBAT thermogenesis (Figure 4E), simulating the cold-stimulated response typically observed in adult mice exposed to 4°C. As expected, 5-day-old (P5) *Plvap^−/−^* mice’s body temperature was significantly lower than their WT littermates (Figure 4F). Taken together, these results suggest that the deficiency in fetal liver-derived macrophages may cause alterations in early BAT function.

## Discussion

Here we show that fetal liver-derived resident macrophages are closely associated with developing sympathetic neural networks in iBAT, and deficiency of these macrophages correlates with dysfunction of the iBAT thermogenesis. BAT is responsible for non-shivering thermogenesis activated in a sympathetic nervous system-dependent mechanism ^50–52^. Non-shivering thermogenesis is a crucial function during the neonatal period, as the adaptation to an out-of-uterine (cold) environment requires heat production to maintain body temperature ^53^. Although iBAT forms already in utero, it continues to develop during postnatal growth and matures from birth until puberty. During maturation, BAT undergoes a thermogenic activity surge starting from P0, which peaks at P14 ^16^. Understanding these stages of target tissue development is crucial, as functional BAT in adults, including humans, plays a key role in energy dissipation and regulation of triglyceride and glucose metabolism ^2–4^. Thus, BAT is a highly interesting tissue for treating obesity and improving glycemic control ^5–8^. Identifying iBAT immune cell content and elucidating the underlying mechanisms controlling pre- and postnatal iBAT growth and stimulation of thermogenic adipocytes could have important implications in developing brown fat-based therapeutics.

During fetal development, progenitors from the yolk sac and the fetal liver give rise to tissue-resident macrophages, discretely populating most developing organs ^47,54–56^. Due to diverse origins and the influence of tissue-specific microenvironments, tissue-resident macrophages are heterogeneous and exhibit vital tissue-specific functions during development and adulthood ^57–59^. Although bone marrow-derived macrophages have been previously identified as the main contributor for BAT macrophages in adult mice ^19^, the present study uncovered the coexistence of several macrophage subsets already in steady-state developing iBAT. We identified F4/80^Int^ macrophages as early as E15.5, simultaneously with the formation of iBAT depots ^9,14^. As the tissue expands and develops, the number of macrophages grows until 1 week of age and then steadily decreases. Interestingly, this decrease is associated with the time when the origin of iBAT macrophage pool is changing as the bone marrow-derived macrophages seed the tissue and eventually substitute the embryonic-derived counterparts in iBAT.

The signature of early iBAT macrophages is very monocytic-like. The flow cytometric analyses revealed that 50-80 % of F4/80^Int^ macrophages during development are positive for Ly6C. CX3CR1 has been previously associated with BAT resident macrophages in adult mice as sympathetic nerve-associated macrophages ^60^ and local regulators of norepinephrine ^18^. From our data, we could detect CX3CR1-positivity in roughly half of the F4/80^Int^ macrophages during iBAT development. The scRNA studies of the clusters identified as iBAT macrophages according to *Fgcr1*, *Adgre1,* and *Csf1r* expression also showed a monocytic signature with the expression of *Ly6c2* and *Ccr2*. *Cx3cr1* and *Trem2* were also expressed in many of the cell clusters. Overall, every iBAT macrophage cluster shared a similar core RNA expression pattern, suggesting the existence of somewhat different cell fates rather than diverse cell populations. Interestingly, *Mrc1*, gene coding CD206, was identified only in one small cluster expressing *Pf4* and *Lyve1*. In mature BAT, CD206 macrophages have been described as a prominent cluster of tissue-resident cells, which increase upon cold exposure or sympathetic nervous system (SNS) stimulation ^19,61,62^. The absence of CD206^+^ macrophages in prenatal iBAT argues for a possible difference in macrophage function at this stage of development. Furthermore, antigen-presenting genes (e.g., *H2-Ab1* and *H2-Eb1*) were identified only in a small population in embryonic and adult iBAT at all studied time points, also suggesting that iBAT macrophages may serve functions beyond their canonical role.

The monocytic signature of iBAT macrophages, both in proteomic and transcriptomic levels, indicates the fetal liver origin in prenatal mice and bone marrow origin in postnatal mice. Genetic fate mapping analyses verified the fetal liver as the primary source of tissue-resident macrophages in developing iBAT and verified that the yolk sac does not contribute to iBAT macrophages. However, soon after birth, at P14, fetal liver-derived cells had already diminished in iBAT and were utterly absent from the iBAT of P35 mice data not shown. The *Ms4a3^CreERT^*^2^*; R26^TdTomato^* model, targeting bone marrow-derived cells ^42^, revealed that newly generated bone marrow-derived monocytes are seeding the tissue already at 1 week of age, and they substitute entirely fetal liver-derived macrophages by P35 data not shown. These data are in line with Gallerand et al. showing that bone marrow is the main source for iBAT macrophages in adult mice ^19^.

There is clear evidence of tissue-resident macrophages controlling the activity of BAT by cross-interacting with nerve fibers and brown adipocytes in adult mice ^17,18,60–62^. Macrophages in mature iBAT have shown to be connected with sympathetic nerve fibers, maintaining the innervation of the tissue ^18^ and mediating the clearance of norepinephrine (NE) via NE transporter, solute carrier family 6 member 2, and degradation enzyme, monoamine oxidase A ^60^. However, the murine sympathetic nervous system is not active in utero and is still developing after birth, reaching maturation only by puberty ^9,13,14,63,64^. Interestingly, we observed that embryonic, fetal liver-derived macrophages were already preferentially organized in clusters closely interacting with developing sympathetic neurons, while co-localization with blood vessels was not seen. Our data indicate a synchronized timeline between the development and maturation of iBAT and the appearance of embryonic macrophages, supporting a possibility that embryonic macrophages may serve an essential functional role in the maturation of sympathetic nerve fibers. However, it appears that the presence of macrophages is not necessary for the normal morphological formation of the nerve network of sympathetic nerve fibers. Our data of *Plvap* deficient mice, lacking fetal liver-derived macrophages but having a normal contribution of the yolk sac and bone marrow-derived cells, further supported this concept. The relative iBAT weight was not altered in postnatal *Plvap^−/−^* mice compared to WT littermates. Nonetheless, iBAT development was delayed in prenatal *Plvap^−/−^* embryos, as morphology at E17.5 was more comparable to WT E15.5 than E17.5 timepoint. Furthermore, postnatal iBAT histology showed similar tissue architecture for *Plvap^−/−^*and WT littermates, and the sympathetic nerves were comparable to WT controls when analyzed by TH whole mount staining from pre- and postnatal iBAT. Although, having a normal morphological tissue development, bulk RNA sequencing analyses of iBAT of P14 *Plvap^−/−^* mice showed significant downregulation of genes associated with thermogenesis and PPAR signaling pathway, indicating a deficiency in the metabolic response and differentiation of brown adipocytes in the absence of PLVAP. Taken together, the cell fate mapping studies and localization of fetal liver-derived macrophages in developing iBAT suggest that fetal liver macrophages have a role in supporting the development and maturation of sympathetic nerve fibers, and after the neural networks are established, fetal liver-derived macrophages are entirely replaced with bone marrow-derived macrophages.

BAT is a critical thermogenic organ that helps mammals to survive in the cold. Human infants and small animals have a high surface area-to-volume ratio losing more heat than adults to the environment. BAT plays a vital role in the adaptation to the out-of-uterine environment after birth. Exposure to a colder environment triggers the SNS activation and noradrenaline release by SNS fibers in BAT, leading to non-shivering thermogenesis. This process is essential for maintaining a stable body temperature in infants as the muscles have not yet developed their thermogenic shivering function^53^. Here we show that the absence of fetal liver-derived macrophages correlated to impaired adaptation to cold in newborn mice, significantly lowering the iBAT temperature (thermogenesis) in neonatal *Plvap^−/−^* mice compared to their WT littermates. Further studies to reveal the role of macrophages in early-life iBAT SNS function are still needed. Interestingly, depleting BAT macrophages in adult mice using CD169^DTR^ model also resulted in impairment in thermogenic function ^61^.

We identified iBAT macrophages as fetal liver-derived during development and their replacement in early postnatal life by bone marrow-derived macrophages. Fetal liver-derived tissue-resident macrophages closely interacted with the developing neural network, possibly suggesting a function in synapse development and neuron maturation. Indeed, the lack of fetal liver-derived macrophages associated with a defect in early-life thermogenesis. Our data suggest that fetal liver-derived macrophages interacting with iBAT’s developing sympathetic neural network may regulate early-life iBAT thermogenesis in mice. Identifying brown adipose tissue immune cell content and elucidating the underlying mechanisms controlling pre- and postnatal iBAT growth and stimulation of thermogenic adipocytes could have important implications in developing brown fat-based therapeutics.

## Methods

### Mice

Multiple genetic mouse models were used in this study: *Plvaptm1Salm* mice (referred to as *Plvap^−/−^*) have been described previously ^22,65^. *R26^EYYP^* (006148), *R26^TdTomato^* (007914), and *Cx3cr1^CreERT2^* (020940) were purchased from Jackson Laboratories. *CCR2^creERT2BB^* mice were kindly provided by Prof. Dr. Burkhard Becker, and *Ms4a3^CreERT2^* mice were kindly provided by Prof. Florent Ginhoux. Wild type (WT) mice, C57BL/6N, were acquired from Janvier labs. All experiments were conducted following the rules and regulations of the Finnish Act on Animal Experimentation (497/2013) and according to the 3R principle under Animal License ESAVI/14685/2020. All experimental mice were housed at 22°C and 12 hours of light and 12 hours of dark cycles at the animal facilities of the University of Turku (Turku, Finland). Standard mouse chow diet and tap water were provided ad libitum. Embryonic development was estimated considering the day of a vaginal plug as embryonic day 0.5 (E0.5).

### In utero and adult tamoxifen administration

*Cx3cr1^CreERT2^* and *CCR2^creERT2BB^* mice were crossed with *R26^EYFP^* or *R26^TdTomato^* reporter mice to study embryonic-derived macrophages. The *Ms4a3^CreERT2^* fate-mapping mouse model was crossed with the *R26^TdTomato^* reporter strain to study bone marrow-derived macrophages. For in-utero tamoxifen induction of CRE activity, a single dose of tamoxifen (1.5 mg, mended with 0.75 mg progesterone; Sigma–Aldrich) was administered i.p. to pregnant females. The E9.5 induction of *Cx3cr1^CreERT^*^2^*; R26^EYFP^* labels yolk sac-derived macrophages only since CX3CR1 is not expressed in fetal liver-derived monocytes or their precursors ^32^. For postnatal tamoxifen induction, *Ms4a3^CreERT2^; R26^TdTomato^* mice were administered i.p., a single dose of tamoxifen (0.56 mg per dose). This induction scheme leads to the selective labeling of MS4A3-positive bone marrow-derived monocytes ^42^.

### Cytometry

To gain a single-cell suspension, iBAT was collected to Hanks’ Buffered Saline, minced with scissors, and incubated with enzymes (1 mg/ml collagenase D and 50 µg/ml DNase 1) for 1h at 37 ℃. After digestion, ≥5-week-old iBAT required gravitational separation of stromal vascular fraction (SVF), filtering through silk (77µm), washing, and suspension in PBS before staining with fluorescent-tagged antibodies for flow cytometric and cell sorting analyses. The list of antibodies and reagents used are listed in the key resources table. Control tissues were dissociated into a single-cell suspension with different procedures. Briefly, blood was collected by cutting the head and draining the blood to 500µL EPICS I with 4mM EDTA (<2-week-old mice) or through cardiac puncture with a heparin-coated syringe (≥2-week-old mice). Erythrocytes were lysed by hypotonic lysis. Brains were dissociated for 30 min at 37 ℃, and microglia were isolated, followed by a Percoll gradient. Lung was mechanically dissociated with scissors, followed by enzymatic digestion for 1 hour at 37℃, filtered through a 70 µm cell strainer, and silk.

Single-cell suspensions were incubated with eFluor780 viability dye for 30 minutes for flow cytometry and sorting. Consequently, Fc-receptors were blocked with anti-CD16/32 for 10 minutes. A fluorochrome-conjugated antibody mix was added to the samples and incubated for 20-30 minutes. Samples were freshly acquired or fixed in a 2 % PFA buffer until acquisition. Flow cytometry data were acquired with an LSR Fortessa flow cytometer (Becton Dickinson) and analyzed using the FlowJo software (FlowJo LLC). Cell sorting was performed in Sony SH800 Cell Sorter (Sony Biotechnology Inc.); the single (FSC-H versus FCS-W plot and SSC-H versus SSC-W) live (eFluor780^−^) CD45^+^ cells were collected to RPMI medium with 2 % FCS.

### Single-cell RNA sequencing

Single live CD45^+^ freshly sorted cells from iBAT at E17.5, P0, and P7 animals were immediately processed according to 10X Genomics guidelines (CG000126_Guidelines for Optimal Sample Prep Flow Chart RevA). Single-cell RNA-sequencing libraries were prepared using Chromium Next GEM Single Cell 3ʹ Kit v3.1 (PN-1000268), Chromium Next GEM Chip G Single Cell Kit (PN-1000120), and Dual Index Kit TT Set A (PN-1000215) according to Chromium Next GEM Single Cell 3’ Reagent Kit, Dual Index v3.1 User Guide (CG000315, Rev A). P0 BAT library was sequenced using Illumina’s Hiseq3000 sequencer. E17.5 and P7 BAT libraries were sequenced using Illumina’s Novaseq6000. Each sample was sequenced to an average depth of ∼50 000 reads per cell. Post-processing, including demultiplexing, read alignment, and quality control, was performed using the 10X Genomics Cell Ranger package (v2.0.0 or v2.1.1).

Pre-processed data were analyzed by Seurat (v4.3) for graph-based clustering and analysis of differentially expressed genes. For quality control, we removed genes expressed in less than 3 cells and cells which expressed less than 200 genes. We also filtered out cells with more than 5000 unique gene counts and 10 % mitochondrial genes. Data were normalized by the function of “NormalizeData,” and variable genes across the single cells were detected by the FindVariableFeatures function. To reduce the influence of variability in the number of UMIs, mitochondrial gene expression, and ribosomal gene expression between cells, the ScaleData function was used. The cells from 3 samples were integrated, and FindClusters was used to apply shared nearest neighbor graph-based clustering to the combined cells using the resolution 0.8. The clustering was visualized with Uniform Manifold Approximation and Projection (UMAP) using RunUMAP and DimPlot.

Differentially expressed genes between the clusters were calculated with the function FindAllMarkers of Seurat (ver 4.3). Min.pct (minimum percentage of the gene-expressing cells in either of all clusters) was set to 0.25 (25%) and thresh.use (minimum fold change in the gene expression between each cluster to all other clusters) was set to 0.25 (log2FC). Violin plots, dot plots, and heatmaps were used to visualize expression by the VlnPlot, DotPlot, and DoHeatmap functions, respectively. Clusters expressing macrophage signature were subclustered by using resolution 0.7. Differentially expressed genes were analyzed as described above.

### Histology

For histological analyses, iBAT was collected and fixed with 10 % buffered formalin solution at RT for 24h. Five µm thick sections were cut, deparaffinized and rehydrated, and stained with the Periodic acid-Schiff (PAS) staining method. The sections were imaged using a Pannoramic 1000 slide scanner (3DHistec).

### Wholemount immunofluorescence

iBAT was collected to PBS and cut with scissors into ≈0.3 mm2 pieces. Briefly, the tissue was fixed overnight at 4 °C in IC (intracellular) Fixation buffer (Thermo Fisher Scientific) diluted 1 in 4 with PBS. Blocking was performed with 1 % species-specific sera according to the antibody mix used, 0.3 % Triton X-100 and 1 % FCS. Primary antibodies were incubated for 3 days at 37 °C, secondary antibodies for 24-32 hours with one day wash with blocking buffer in between. After washing, in washing buffer (PBS + 0.2 % Triton X-100 + 0.5 % 1-thioglycerol), the samples were dehydrated at 4 °C in isopropanol in H_2_O gradient 30-100 % for 3h, bleached for 2h (isopropanol + 5% DMSO + 2% H_2_O_2_) and incubated 1h in 100 % isopropanol. Before image acquisition, the samples were stored overnight in a clearing medium (ECi, ethyl cinnamate). Image acquisition, processing, and analysis was performed with 3i CSU-W1 spinning disk confocal microscope (Intelligent Imaging Innovations) equipped with Photometrics Prime BSI sCMOS camera (Teledyne Imaging) and Plan-Apochromat Zeiss 10x/0.45 or Plan-Apochromat Zeiss 20x/0.8 dry objectives, using SlideBook 6 software (Intelligent Imaging Innovations) and ImageJ software (1.54b). The lasers used were 488 nm, 561 nm, and 640 nm, and the filters used were GFP (525/30nm), Cy3/Alexa568 (617/73nm), and Cy5/Alexa647 (692/40nm).

### Bulk RNA sequencing

Whole iBAT from P14 *Plvap^−/−^* mice and control littermates was collected to RNA later and stored at −20 °C until RNA isolation with Macherey-Nagel Nucleospin RNA kit and eluted in RNase-free water. Isolated total RNA was stored at −70 °C. The quality of the samples was ensured using Agilent Bioanalyzer 2100, and their concentration was measured with Qubit®/Quant-IT® Fluorometric Quantitation, Life Technologies, before and after library preparation. Library preparation was performed at the Finnish Functional Genomics Centre, University of Turku and Åbo Akademi and Biocenter Finland with Illumina Stranded mRNA Preparation, Ligation, Kit (Illumina), according to Illumina Stranded mRNA Library Preparation, Illumina (1000000124518) protocol. Next-generation sequencing was performed using Illumina NovaSeq 6000 instrument.

Raw reads were aligned to mouse genome build GRCm38 using HISAT2 alignment program^66^. The gene expression level was estimated by the abundance of transcripts (count of sequencing) mapped to the genome or exon. Read counts were proportioned to gene expression level, length, and sequencing depth. FPKM (expected number of Fragments Per Kilobase of transcript sequence per Millions of base pairs sequenced) method was used to estimate gene expression levels ^66^. We performed Principal Component Analysis (PCA) analysis on the gene expression value (FPKM) of all samples after filtering and normalizing (CMP: counts per millions) the data. Differentially expressed genes were then analyzed using DESeq2 (v1.36.0) package. Gene enrichment analyses of the differentially expressed genes were performed using Gene Ontology (GO, http://www.geneontology.org/) and the Kyoto Encyclopedia of Genes and Genomes (KEGG).

### Thermoneutrality experiment

Timed matings were set at RT, and at E11.5±2 days, females identified as pregnant were randomly assigned to two separate cohorts: pregnant dams housed at RT vs. pregnant dams housed at 30 ℃. The iBAT from the offspring of the two cohorts was collected at P0, P7, and P14. Samples were analyzed by flow cytometry.

### Infrared thermography

Offspring from heterozygous *Plvap* breeding were analyzed at 5 days of age (P5). The pups were separated from their huddle, individually kept with separators in an area ranging from 49-92 cm^2^ for 30 minutes at RT (acclimatization), and then photographed with FLIR T620bx (Infradex oy) thermal camera. Genotyping was performed at P10 to identify the *Plvap^−/−^* and control littermates *Plvap^+/+^*. The infrared photos were analyzed, and the individual interscapular maximum temperature for each individual pup was acquired using FLIR Thermal Studio – 1.9.40.0 software from FLIR systems.

### Quantification and statistical analysis

Numeric data are given as mean ± SEM. GraphPad Prism (v9 GraphPad Software, LLC) was used for the statistical analyses. Comparisons between the time points, genotypes, or treatment groups were made using the nonparametric two-tailed Mann-Whitney test, parametric two-tailed t-test, and one- or two-way ANOVA test with Bonferroni post-hoc test. P-values lower than 0.5 were considered statistically significant.

## Supplemental information

**Supplementary Figure 1.**
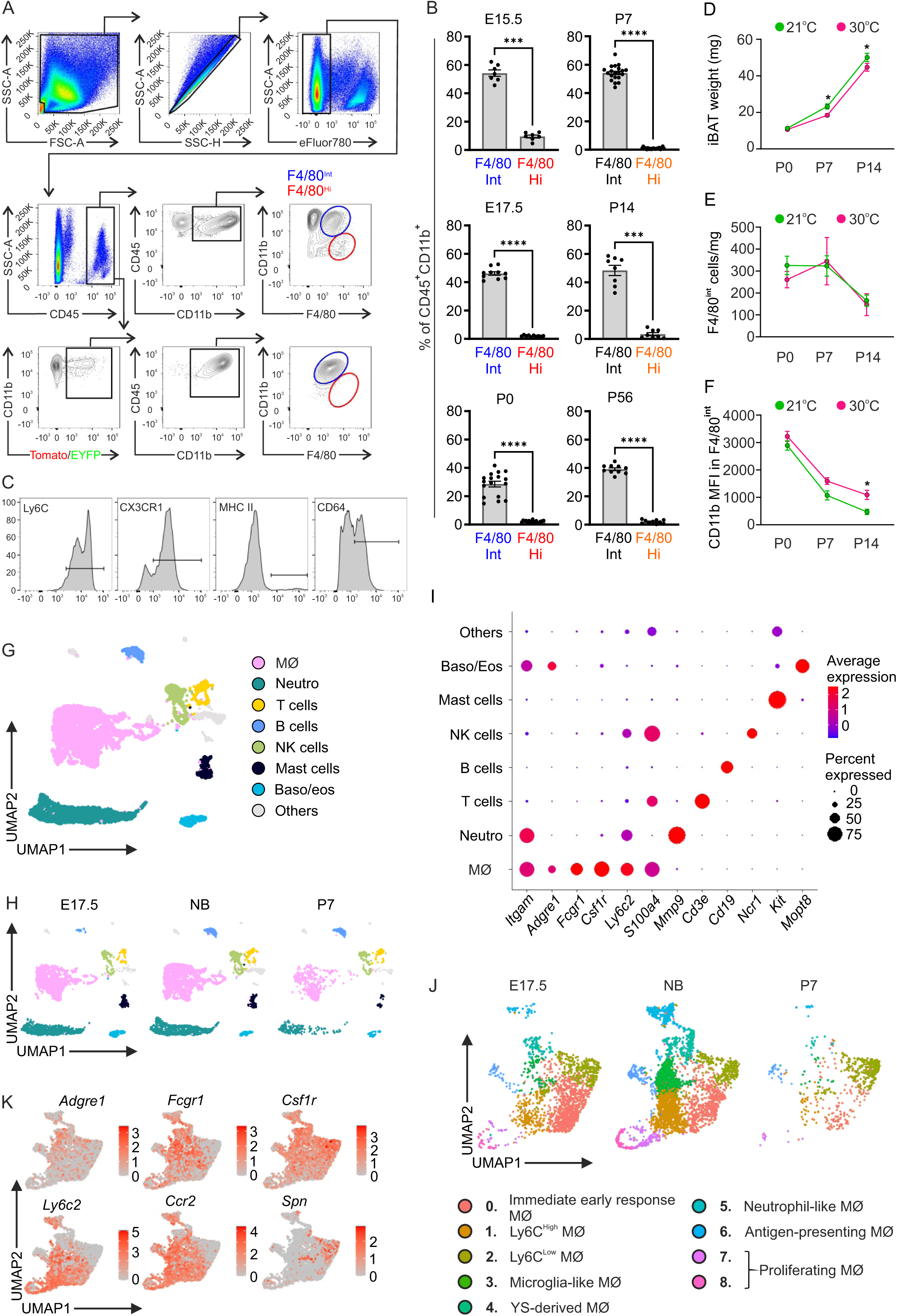
The diversity of leukocyte and macrophage subsets in early iBAT. (A) Gating strategy of the flow cytometric data for iBAT single live (eFluor780^−^) CD45^+^CD11b^+^F4/80^Int/Hi^ macrophages and for TdTomato/YFP positive cells. (B) Frequencies of F4/80^Int^ and F4/80^Hi^ cells in iBAT at indicated time points (n = 7-20/time point). (C) Representative histograms of Ly6C, CX3CR1, MHC II, and CD64 surface markers in F4/80^Int^ cell population. (D) iBAT weight of WT mice at RT (21 ℃) and thermoneutral environment (30 ℃) at indicated time points. (E) F4/80^Int^ cell numbers related to iBAT weight (mg) of WT mice at RT (21 ℃) and thermoneutral environment (30 ℃) at the indicated time points. (F) Mean fluorescence intensity (MFI) of CD11b in F4/80^Int^ population of WT mice at RT (21 ℃) and thermoneutral environment (30 ℃) at indicated time points. (G) UMAP visualization of CD45^+^ leukocytes in iBAT of E17.5, P0, and P7 mice combined. MØ, macrophages; Neutro, neutrophils; Baso/eos, basophils and eosinophils. (H) UMAP visualization of CD45^+^ leukocytes in iBAT of E17.5, P0, and P7 mice individually. (I) Average gene expression dot plot of genes identified in main BAT leukocyte types. (J) UMAP visualization of macrophages expressing *Adgre1*, *Fcgr1*, and *Csf1r* in iBAT of E17.5, P0, and P7 mice separately. (K) Gene expression of selected genes on top of UMAP visualization of macrophages in iBAT of E17.5, P0, and P7 mice combined.

**Supplementary Figure 2.**
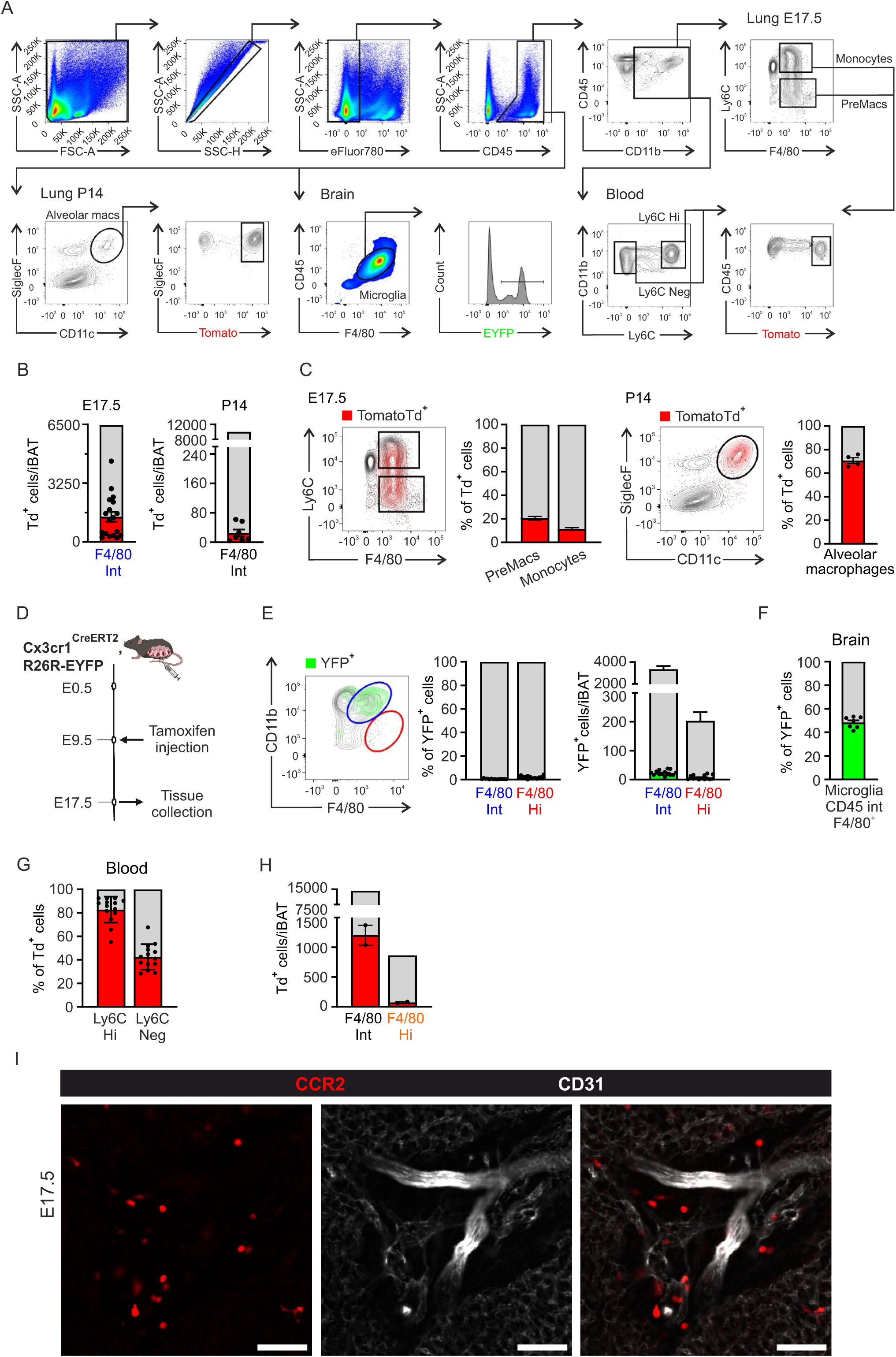
Early iBAT macrophages originate from the fetal liver and are replaced by the bone marrow-derived progenitors in juvenile mice. (A) Gating strategy of the flow cytometric data for lung, brain and blood single live (eFluor780^−^) CD45^+^ cells. (B) TdTomato^+^ cell numbers per iBAT of *Ccr2^CreERT^*^2^*; TdTomato* mice in F4/80^Int^ at indicated time points. Mean±SEM. (C) Frequency of TdTomato^+^ cells in lung of *Ccr2^CreERT^*^2^*; TdTomato* mice at indicated time points. Mean±SEM. (D) Experimental setup of the induction scheme of *Cx3cr1^CreERT^*^2^*; R26EYFP* mice. (E) Representative flow cytometry plot, frequency of YFP^+^ cells, and YFP^+^ cell amounts in F4/80^Int^ and F4/80^Hi^ macrophages per iBAT of E17.5 *Cx3cr1^CreERT^*^2^*; R26EYFP* mice. Mean±SEM. (F) Frequency of YFP^+^ cells in CD45^Int^F4/80^+^ microglia of E17.5 *Cx3cr1^CreERT^*^2^*; R26EYFP* mice. Mean±SEM. (G) Frequency of TdTomato^+^ cells in Ly6C^Hi^ and Ly6C^Neg^ cells in blood of *Ms4a3^CreERT^*^2^*; R26^TdTomato^*mice. Mean±SEM. (H) TdTomato^+^ cell amounts in F4/80^Int^ and F4/80^Hi^ macrophages per iBAT of P7 *Ms4a3^CreERT^*^2^*; R26^TdTomato^* mice. Mean±SEM. (I) Wholemount immunofluorescence staining of *Ccr2^CreERT^; TdTomato* mice iBAT at E17.5 with anti-CD31 (white) and anti-RFP (red) detecting TdTomato. Bar representative of 50 µm.

**Supplementary Figure 3.**
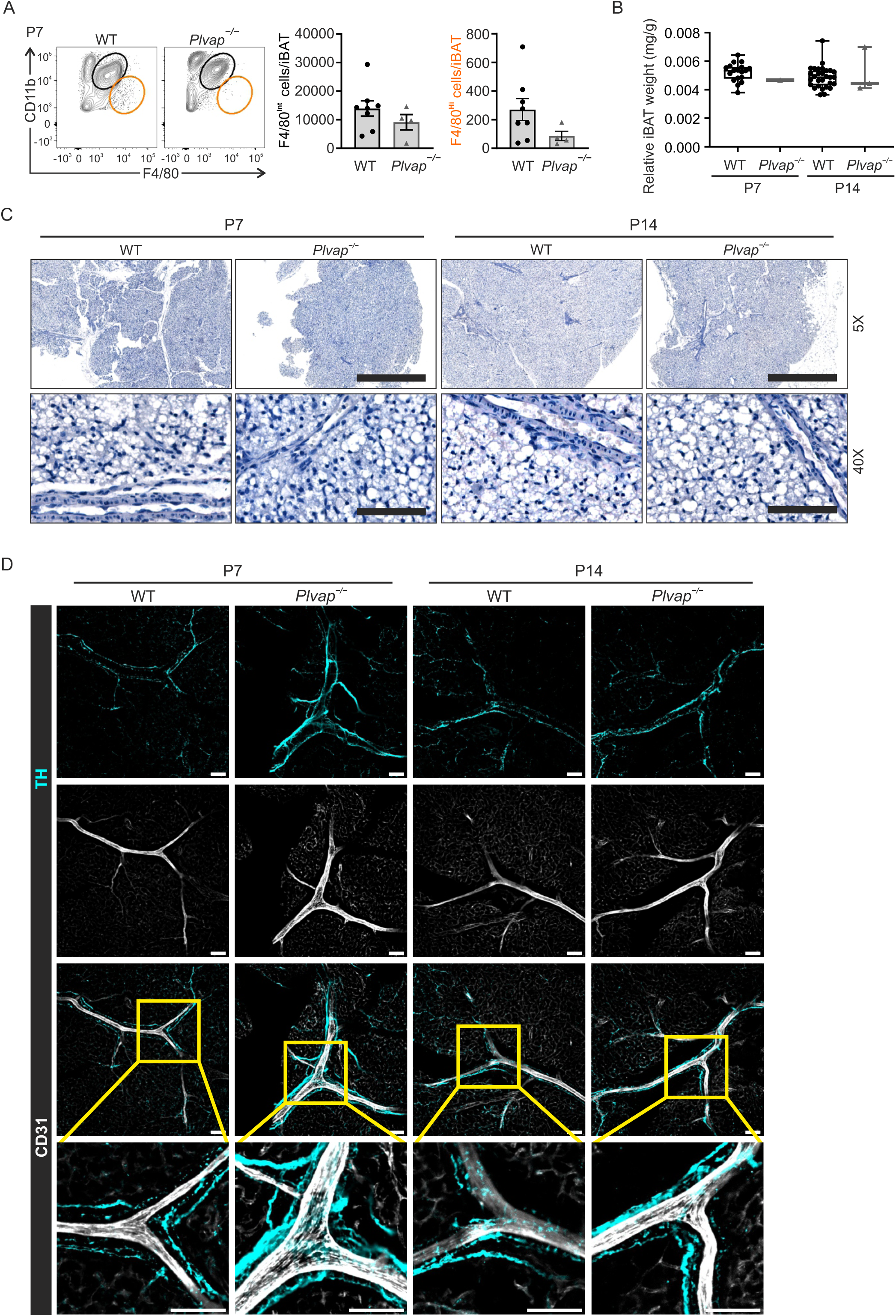
The postnatal development of iBAT in *Plvap^−/−^* mice lacking fetal liver-derived macrophages is normal. (A) Representative flow cytometry plots and cell numbers of F4/80^Int^ and F4/80^Hi^ macrophages per iBAT of *Plvap^−/−^* and WT mice at P7 (n = 4-8). Mean±SEM. (B) iBAT weight relative to bodyweight of *Plvap^−/−^* compared with WT mice at P7 and P14. Mean±SEM. (C) *Plvap^−/−^* iBAT PAS stained histology compared with WT mice at P7 and P14. Scale bars are 50 µm in overview (5X) and 500 µm in 40X images. (D) Wholemount immunofluorescence staining of *Plvap^−/−^* mice iBAT at P7 and P14 with anti-TH (cyan) and anti-CD31 (white). Scale bars are 50 µm.

**Supplementary Figure 4.**
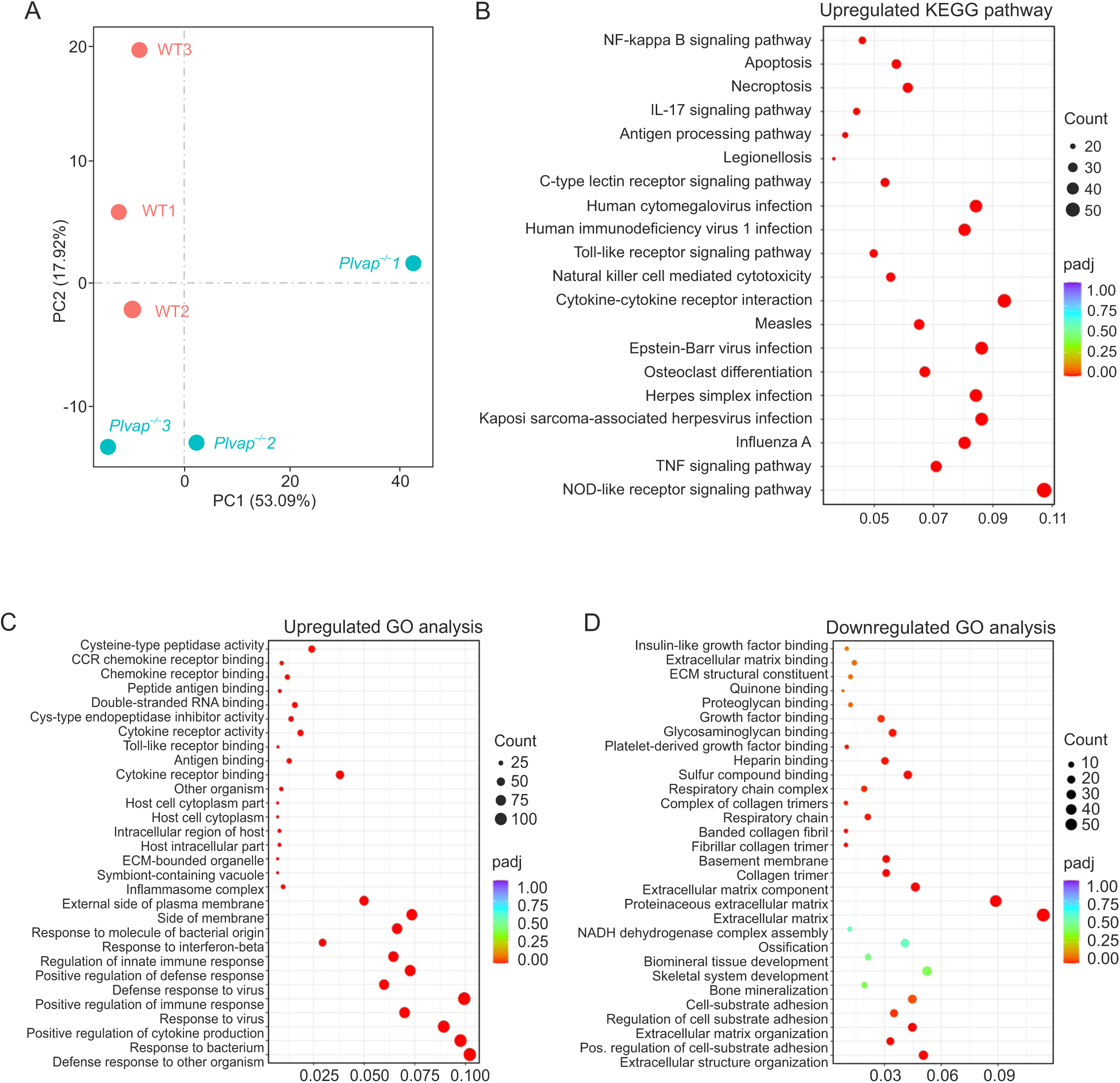
Macrophage deficiency during development associates with early BAT dysfunction. (A) Bulk RNA sequencing principle component analysis (PCA) of gene expression in iBAT of P14 *Plvap^−/−^* compared with WT controls (n = 3). (B) KEGG enrichment analysis of the top 20 upregulated pathways in iBAT of *Plvap^−/−^*. (C) GO annotation analysis of the top 20 upregulated genes in iBAT of *Plvap^−/−^*. (D) GO annotation analysis of the top 20 downregulated genes in iBAT of *Plvap^−/−^*.

**Supplementary Table 1.**
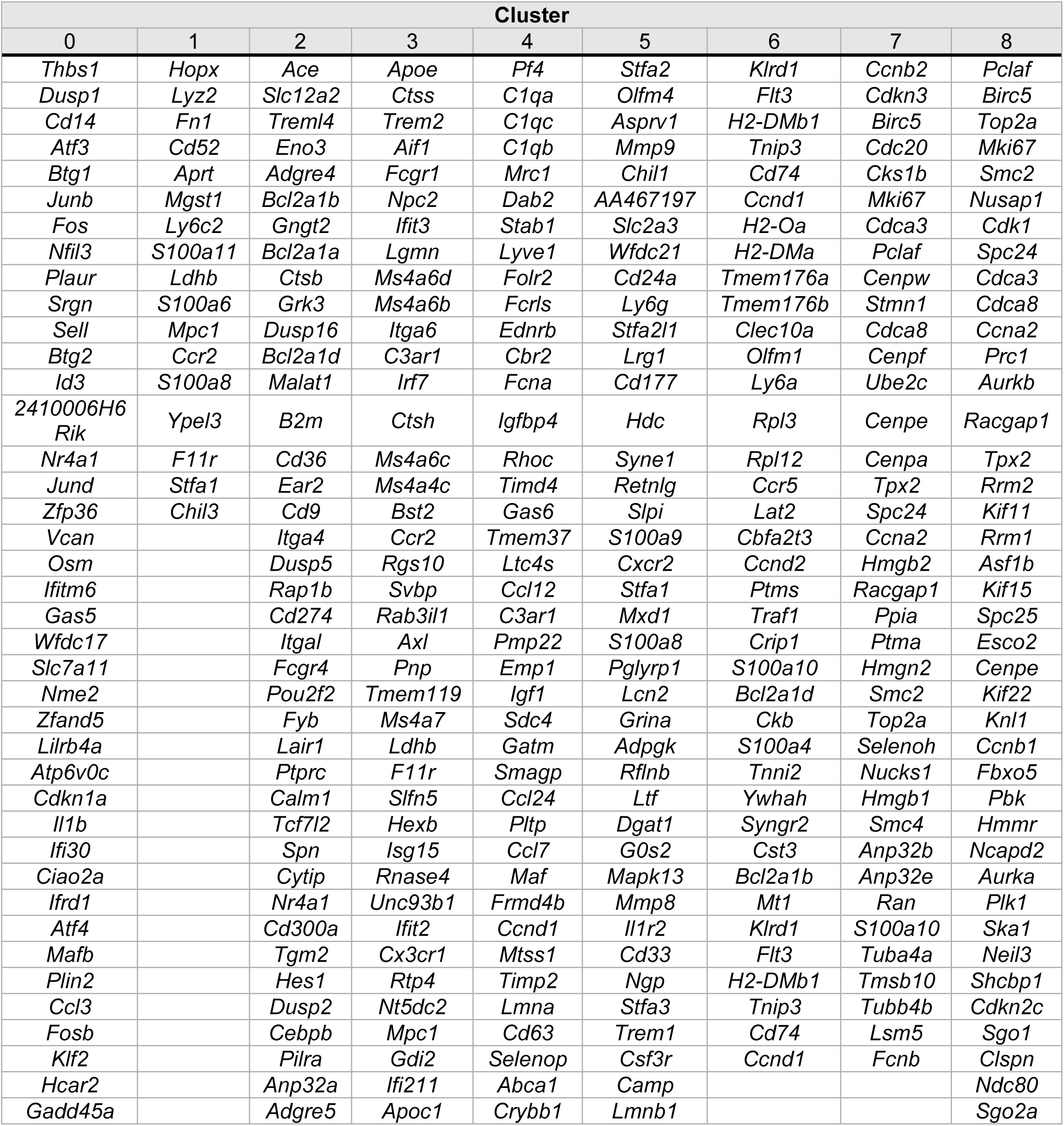
Top 40 upregulated genes of each macrophage cluster organized by adjusted p-value.

